# Environmental drivers of heritable trait variation and lag of adaptation to climate in *Hordeum murinum*

**DOI:** 10.64898/2026.08.11.744122

**Authors:** Helene Villhauer, Sandy Jan Labarosa, Timo Hellwig, Sabine Ambrosius, Przemyslaw Baranow, Albane Bignon, José Manuel Blanco Moreno, Denise V. M. Blume, Anna Bomanowska, Milan Brankov, Nina Döring, Walter Durka, Severin Einspanier, Arndt Hampe, Miloš Ilić, Konrad Kaczmarek, Abdenour Kheloufi, Lea Klepka, Marta Kolanowska, Kamil Konowalik, Stanislav Kopriva, Izabela Krzemińska, Mathieu Leclerc, Lena Lerbs, Sascha Liepelt, Lahouaria Mounia Mansouri, Víctor Manzanares-Vázquez, Sabine Metzger, Nadine Mitschunas, Monika Myśliwy, Pablo Neira, Agnieszka Nobis, Marcin Nobis, Artur Nosalewicz, Sławomir Nowak, Sylvain Pincebourde, Boris Radak, Agnieszka Rewicz, Encarna Rodríguez-García, Aritz Royo-Esnal, Francesco Santi, Luís P. da Silva, Henryk Straube, Jannis Straube, Katalin Szitár, Joel Torra, Markus Wagner, Adrian Wysocki, Maria von Korff, Anna Bucharova

**Affiliations:** Conservation Biology, Department of Biology, Philipps University Marburg, 35043 Marburg, Germany; Institute of Plant Genetics, Department of Biology, Heinrich-Heine-University Düsseldorf, 40225 Düsseldorf, Germany; Institute for Plant Sciences, University of Cologne, 50674 Cologne, Germany; Department of Plant Taxonomy and Nature Conservation, Faculty of Biology, University of Gdańsk, Wita Stwosza 59, 80-308 Gdańsk, Poland; Laboratoire de Biométrie et Biologie Evolutive (LBBE), UMR 5558, CNRS, Université Claude Bernard Lyon 1, 69100 Villeurbanne, France; Departament de Biologia Evolutiva, Ecologia i Ciències Ambientals, Facultat de Biologia, Universitat de Barcelona (UB), Av. Diagonal 643, Barcelona 08028, Spain; Institut de Recerca de la Biodiversitat (IRBio), Universitat de Barcelona (UB), Barcelona 08028, Spain; Faculty of Science, Department of Plant and Environmental Sciences, Section for Plant Biochemistry, University of Copenhagen, Frederiksberg 1871, Denmark; Department of Geobotany and Plant Ecology, Faculty of Biology and Environmental Protection, University of Lodz, Banacha 12/16, 90-237 Lodz, Poland; Maize Research Institute Zemun Polje, Slobodana Bajića 1, 11185 Belgrade, Serbia; Department of Community Ecology (BZF), Helmholtz Centre for Environmental Research - UFZ, 06120 Halle, Germany; German Centre for Integrative Biodiversity Research (iDiv) Halle-Jena-Leipzig, 04103 Leipzig, Germany; Department of Phytopathology and Crop Protection, Institute of Phytopathology, Faculty of Agricultural and Nutritional Sciences, Kiel University, 2408 Kiel, Germany; INRAE, University of Bordeaux, BIOGECO, F-33610, Cestas, France; Department of Biology and Ecology, Faculty of Sciences, University of Novi Sad, Trg Dositeja Obradovića 2, 21000 Novi Sad, Serbia; Department of Ecology and Environment, University of Batna 2, Batna 05078, Algeria; Department of Botany and Plant Ecology, Wrocław University of Environmental and Life Sciences, 50-363 Wroclaw, Poland; Cluster of Excellence on Plant Sciences "SMART Plants for Tomorrow’s Needs", 40223 Düsseldorf, Germany; Institute of Agrophysics, Polish Academy of Sciences, Doświadczalna 4, 20-290 Lublin, Poland; Institut de Recherche sur la Biologie de l’Insecte, UMR 7261, CNRS - Université de Tours, 37200 Tours, France; École Supérieure des Agricultures, ESA, 49000 Angers, France; Department of Research and Development, Coccosphere Environmental Analysis, Calle Cruz 39, 29120-Alhaurín el Grande, Málaga, Spain; Institute for Plant Sciences, Biocenter MS-Platform, University of Cologne, 50674 Cologne, Germany; UK Centre for Ecology & Hydrology, Benson Lane, Wallingford, Oxfordshire, OX10 8BB, UK; Department of Environmental Ecology, Institute of Marine and Environmental Sciences, University of Szczecin, Mickiewicza 16, 70-383 Szczecin, Poland; Institute of Botany, Faculty of Biology, Jagiellonian University, Gronostajowa 3, 30-387 Kraków, Poland; Active Learning in Ecology and Biotechnology. Calle Las Moreras, 5. 30149. El Siscar-Santomera-Murcia (Spain); Department of Agricultural and Forest Sciences and Engineering, Universitat de Lleida — Agrotecnio CERCA Centre, Lleida 25198, Spain; Serra Húnter Programme, Universitat de Lleida (UdL), Lleida 25198, Spain; Biome Lab, Department of Biological, Geological and Environmental Sciences, Alma Mater Studiorum - University of Bologna, 40126 Bologna, Italy; EBM, Estação Biológica de Mértola, Praça Luís de Camões, Mértola, 7750-329 Mértola, Portugal; CIBIO, Centro de Investigação em Biodiversidade e Recursos Genéticos, InBIO Laboratório Associado, Campus de Vairão, Universidade do Porto, 4485-661 Vairão, Vila do Conde, Portugal; MTA–HUN-REN Centre for Ecological Research, ‘Lendület’ Landscape and Conservation Ecology Research Group, Institute of Ecology and Botany, 2–4 Alkotmány út, Vácrátót, 2163, Hungary; Department of Plant Biology, Institute of Biology, Wrocław University of Environmental and Life Sciences, Kożuchowska 7a, Wrocław PL-51-631, Poland

**Keywords:** climate adaptation, common garden experiment, grasses, evolutionary lag, trait variation, *Hordeum murinum*, *in situ*, phenotypic plasticity

## Abstract

1. Most plant species are genetically differentiated among populations, often reflected by phenotypic trait variation that corresponds to local adaptation. Yet the strength of local adaptation and heritable contribution to phenotypic traits vary across traits, species, and environments. Additionally, climate change is rapidly altering environmental conditions, and the climate may shift faster than populations can adapt or track the change via dispersal, resulting in adaptive lags. However, it remains unclear how widespread such adaptive lags are across plant species.

2. We focused on *Hordeum murinum*, an annual ruderal grass widespread in Europe. We combined continental-scale *in situ* measurements of 2070 plants across 207 populations with common garden experiments across two contrasting climates and two soil types to disentangle heritable variation from phenotypic plasticity and assess potential adaptive lags under climate change.

3. We found that heritable variation was pronounced in developmental traits, particularly flowering time and plant height, while seed weight, reproductive investment and SLA showed intermediate heritable contribution, and flag leaf area and total biomass were primarily plastic. Heritable trait variation was strongly associated with temperature at the populations’ origin, and trait clines were consistent with *in situ* patterns, suggesting that temperature is the main driver of genetic differentiation in *H. murinum*. However, we detected that fitness peaked in populations originating from warmer climates, indicating that evolutionary responses may not keep pace with rapid environmental shifts.

4. Synthesis: Our results highlight that *H. murinum* harbors substantial heritable variation, shaped primarily by temperature. However, the pace of evolutionary change may be insufficient to track ongoing climate change, leaving populations potentially vulnerable to future environmental conditions.

## Introduction

Plant species are no uniform units, their populations are genetically differentiated (Hartl & Clark, 2007). This genetic differentiation is often reflected in phenotypic trait variation (G effect in quantitative genetic studies) and frequently corresponds to adaptation to local environmental conditions, a phenomenon termed local adaptation (Orr, 2005; Turesson, 1922). Local adaptation arises primarily through natural selection on heritable phenotypic variation, when genotypes with a more advantageous suite of traits in a given environment have higher fitness and thus contribute more to the next generation (Joshi et al., 2001). Accordingly, the response to selection depends on the strength of selection and heritability (Caruso et al., 2020), and thus, the contribution of traits to local adaptation varies across traits, populations, and environments.

In natural environments, observed trait variation does not necessarily reflect heritable adaptation alone but may also result from responses to environmental variation through phenotypic plasticity (E effect in quantitative genetic studies). Phenotypic plasticity describes the ability of a single genotype to produce different phenotypes in response to varying environmental conditions (Valladares et al., 2006). It can affect plant morphology, physiology, or development, and the contribution of plasticity to trait variation depends on traits, populations, and environments (Britton et al., 2026; Kreyling et al., 2019; Matesanz et al., 2020). Growth-related, vegetative traits often show higher plasticity than traits more directly associated with fitness (Stearns & Kawecki, 1994), allowing plants to buffer short-term environmental fluctuations (Kreyling et al., 2019) and maintain performance in heterogeneous environments (Matesanz et al., 2020). In contrast, some reproductive traits like flowering time typically show lower plasticity because they directly determine reproductive success and are therefore strongly constrained by selection (Yan et al., 2021). Consequently, phenotypic plasticity – particularly in vegetative traits – may obscure heritable trait variation in field observations along environmental gradients (Britton et al., 2026; Villellas et al., 2021). Additionally, phenotypic plasticity as a trait can be heritable and vary between populations and genotypes (genotype by environment interaction, GxE in quantitative genetic studies) (Schneider, 2022), but the heritable variation in plasticity is often smaller than genetic variation for trait means (Scheiner, 1993). Accordingly, predicting trait values under natural field conditions remains difficult (but see Villellas et al., 2021).

To disentangle heritable trait variation from phenotypic plasticity, plants are typically grown in common garden experiments (De Villemereuil et al., 2016). Differences among populations that persist in such common environments indicate heritable variation, which can be a consequence of neutral demographic processes or local adaptation to both abiotic and biotic factors (Dorey et al., 2024; Macel et al., 2007). For example, when populations of a species from a large-scale climatic gradient are trialed in a common garden, individuals from warmer and drier areas often grow smaller, flower earlier, and have thicker, denser leaves, resulting in lower SLA (specific leaf area) than their conspecifics from cooler or wetter regions. These patterns are commonly interpreted as adaptations to water limitation and thermal stress (Griffin-Nolan et al., 2025; Kilkenny et al., 2026). At smaller spatial scales, heritable trait variation can be affected by soil conditions (Van Nuland et al., 2019) and biotic interactions such as competition, herbivory, or pollination (Bennett et al., 2016; Dorey et al., 2024)

Understanding the drivers of adaptive trait variation is particularly important in the context of climate change. If adaptation is constrained by factors other than climate, such as soil conditions or biotic interactions (Dorey et al., 2024), plant populations may be limited in their ability to shift their ranges or track suitable environments. A previous study on 1000 grass species identified temperature as the dominant driver of trait variation, whereas soil conditions played a comparatively minor role (Griffin-Nolan et al., 2025). The influence of biotic environments, such as competition, is more variable and species-dependent (Bennett et al., 2016), but they usually impose a smaller selection pressure on trait variation (Caruso et al., 2020) and thus contribute less to plant local adaptation than abiotic factors (Hargreaves et al., 2020). Moreover, the effects of abiotic and biotic factors on heritable trait variation can interact (Dorey et al., 2024). Most studies, however, have focused on heritable variation across either large or small geographic scales and have typically examined abiotic and biotic drivers separately (Briscoe Runquist et al., 2020; Dorey et al., 2024). Therefore, studies that simultaneously consider both abiotic and biotic factors across multiple spatial scales remain scarce.

Climate change is causing shifts in environmental conditions, and plant populations might now face a different environment than the one to which they have adapted in the past. This can result in a lag of local adaptation behind climate change, particularly if the population fails to evolve or track climatic optimum via dispersal. Such mismatches have been experimentally documented for several plant species, e.g., *Arabidopsis thaliana* (Wilczek et al., 2014), *Boecheria stricta* (Anderson & Wadgymar, 2020), and *Trifolium repens* (Albano et al., 2026), and suggested by a meta-analysis of reciprocal transplant experiments (Bontrager et al., 2020). However, this pattern is not consistent across all plant species, e.g., *Festuca estia* (Gonzalo-Turpin & Hazard, 2009), *Polygonum cespitosum* (Sultan, 2003), and *Brassica rapa* (Franks et al., 2007) appear to maintain local adaptation despite climate change. Overall, it remains unclear how common lags in local adaptation to climate change are across plant species (Lambrecht et al., 2007).

In this study, we investigated environmental drivers of adaptive trait variation and its potential lag behind climate change in an annual ruderal grass, *Hordeum murinum*. The species thrives in human-disturbed environments and is dispersed by seeds that adhere to human clothes or animal fur (Davison, 1977). Such effective dispersal could allow the species to migrate and track the climatic optimum of the populations (Krause et al., 2015), and anecdotal evidence indeed suggests that it is spreading in response to climate change (de Groot et al., 1995). To understand the drivers of trait variation and assess the potential lag of local adaptation behind climate change, we measured *in situ* trait variation and local environment for 2070 plants from 207 populations across Europe and Northern Africa, and we collected seeds, which we then grew in two climatically different common gardens under two soil conditions. This design allowed us to quantify the contribution of genetic and environmental factors to trait variation and identify potential environmental drivers of this trait variation. Furthermore, we tested whether the local adaptation of the European *H. murinum* populations keeps pace with climate change. We hypothesize that (i) Variation in reproductive traits is primarily heritable, driven by genetic differentiation (G), while variation in growth-related traits is strongly influenced by environmental conditions, and thus primarily driven by phenotypic plasticity (E). Additionally, the extent of phenotypic plasticity varies among populations (GxE), but the heritable variation for plasticity is smaller than the heritable variation for trait means. (ii) Heritable trait variation is driven primarily by broad-scale climatic gradients and less by local edaphic and biotic factors of the seed origin. Specifically, we expect that plants from warmer regions flower earlier, grow smaller, and have lower specific leaf area (SLA) than plants from colder regions. Additionally, gradients in trait variation observed *in situ* correspond to the gradients observed in the common garden for reproductive (more heritable) but not for growth-related traits (less heritable). (iii) The variation in fitness reflects broad-scale climatic adaptation because the species can effectively disperse, and thus local adaptation can track climate change via dispersal.

## Materials and methods

### Study species

*Hordeum murinum* L. (wall barley) is a winter annual grass, a wild relative of cultivated barley (*H. vulgare* L. subsp*. vulgare*), and is common in most temperate regions. While it is native to the Mediterranean and Europe (Figure 1), it has been introduced to Australia, southern Africa, and North America, where it has become invasive (Davison, 1971). Across most of its range, *H. murinum* is primarily associated with ruderal, human-disturbed habitats with high light availability and low competition (Davison, 1977; Mizianty, 2006; Villhauer et al., 2026). The species typically flowers from early spring to late summer, depending on the region. Flowers are predominantly self-pollinated, and seeds are effectively dispersed through epizoochory. Seeds germinate in late summer or early autumn, and plants overwinter in vegetative state (Davison, 1971; Mizianty, 2006). *H. murinum* is a part of an aggregate taxon *Hordeum murinum* agg. that comprises three subspecies differing by cytotypes (2x, 4x, 6x; Cuadrado et al., 2013). This study focuses on the tetraploid *H. murinum* (2n = 4x = 28, ssp. *murinum*, or *H. murinum* s. str.), because it occurs throughout Europe.

**Figure 1.**
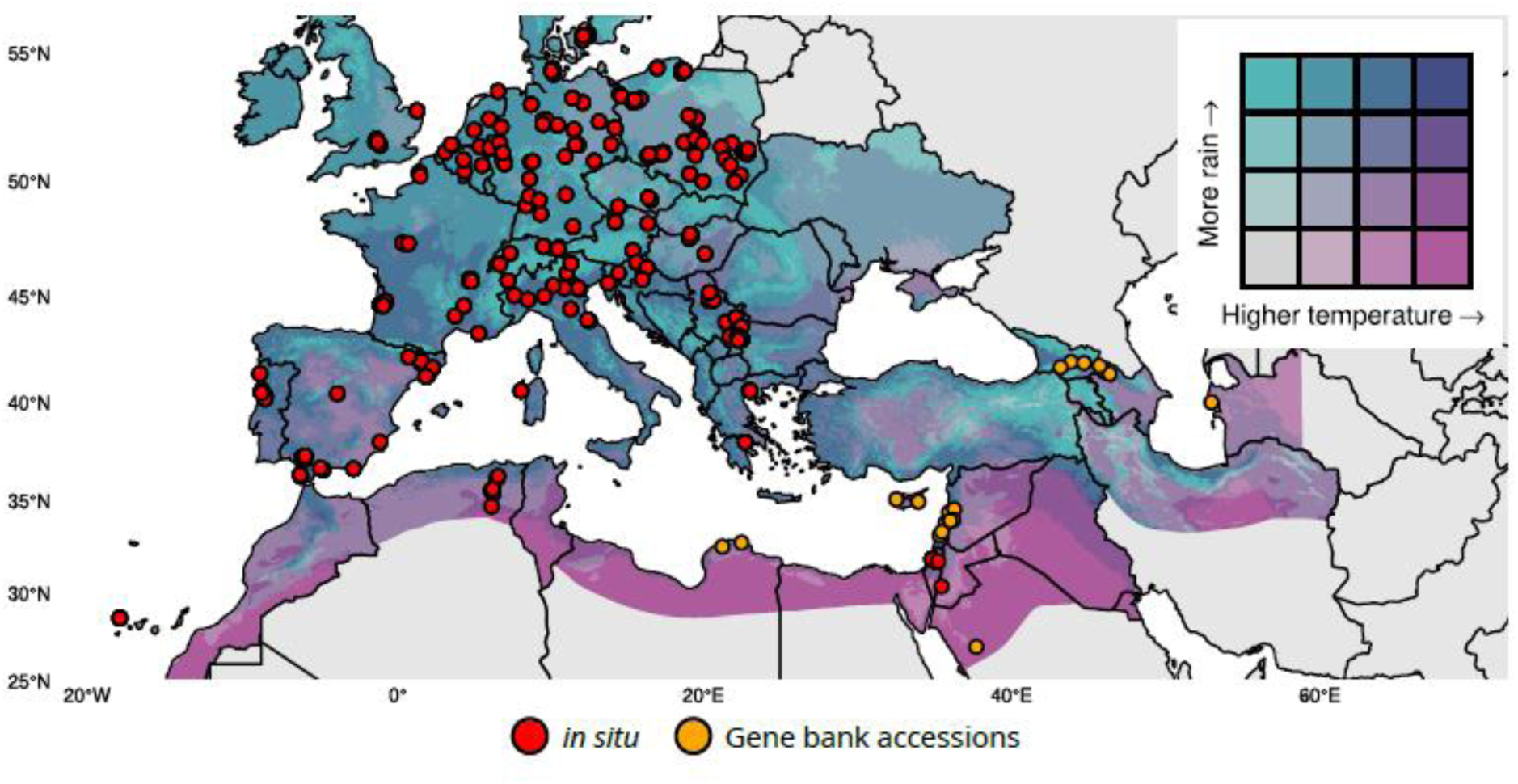
Climate map of the *in situ* sampled populations and gene bank accessions of tetraploid *Hordeum murinum* across Europe, North Africa, and Western and Central Asia. Color shading shows the average climate from the years 2000-2020 of mean annual temperature and annual precipitation.

### Seed collection, in situ traits, environment, and climate

We sampled 207 natural tetraploid populations of *H. murinum* across Europe and neighboring countries between 2022 and 2023 (Figure 1, orange dots, Villhauer et al., 2026). In summary, within each population, we randomly selected 10 plants at least 5 m apart, measured their height, and collected seeds. We considered the seed collection date for each population as a proxy for the onset of seed ripening phenology (March to August, depending on the climate). In the laboratory, we determined seed mass as thousand-seed weight, hereafter referred to as seed weight, and we stored the seeds at 4 °C until further use. To characterize *in situ* biotic and abiotic environmental conditions, we estimated the intensity of competition (for each plant) as the percentage coverage of the surrounding vegetation, and measured soil pH value and bioavailable soil phosphate in the laboratory for each population (Villhauer et al., 2026).

To characterize the climate at seed collection sites, we obtained seven bioclimatic variables for the period 2000–2020 at approximately 1 km² (30 arcsec) spatial resolution from the CHELSA database (Karger et al., 2021). We performed a principal component analysis (PCA) based on these seven variables (Figure S1). Because the first principal component (PC1) was strongly correlated with mean annual temperature (r = 0.90) and the second principal component (PC2) with annual precipitation (r = 0.86), we use mean annual temperature and annual precipitation directly for intuitive interpretation.

To identify the fitness optimum (approximated by reproductive biomass, see below) in relation to temperature, we used mean spring temperature as the climatic predictor, because plants in the common gardens experienced the growing conditions from March to June. We calculated the mean spring temperature as the average of monthly mean temperatures from March to June over the period 2000–2020. As higher-resolution monthly datasets were not available for this period, we extracted monthly temperature at 2.5 arc-minute resolution (∼21 km²) from the WorldClim database (Fick & Hijmans, 2017).

### Common garden experiment

In 2024, we conducted two simultaneous common garden experiments in Düsseldorf (51°11’08.8"N, 6°48’10.2"E, mean spring temperature 13.9 °C in 2024, obtained from Bioclim) and Marburg (50°48’01.9"N, 8°48’24.3"E, mean spring temperature 12.6 °C in 2024, obtained from Bioclim) to assess heritable trait variation and phenotypic plasticity. The two experimental sites experience nearly identical seasonal photoperiods. In Marburg, we grew plants in two growing substrate types: fertilized soil (93% LD80 soil, 7% sand, 250mL Osmocote fertilizer per 70 liters of substrate) and a sand treatment with 70% sand and 30% LD80 soil to simulate low nutrient- and water-holding capacity (further called “sand”). In Düsseldorf, we grew plants only in the soil treatment, using the same substrate mixture as in Marburg. This resulted in three treatments: Düsseldorf soil, Marburg soil, and Marburg sand.

The common garden experiments included seeds from 189 populations. We subsampled the populations to maximise geographic distribution which resulted in a median nearest-neighbor distance of about 24 km. We selected 5 random maternal plants per population and grew one plant per maternal plant (genotype) per treatment. The field-collected seeds were complemented by 19 gene bank accessions to incorporate plants coming from warmer and drier regions (Figure 1, Table S1). For each gene bank accession, we included 4 replicates per treatment. In total, each experimental treatment included 996 plants, summing to 2988 plants across all three treatments.

We initiated the experiment in October 2023 because *H. murinum* is a winter annual that germinates in autumn and requires vernalization to initiate flowering (Davison, 1971). For sowing seeds, we used 12 x 8 plug trays filled with Mini-Tray soil (MIM800, Balster Einheitserdewek, Fröndenberg, Germany) and supplemented with slow-release fertilizer (Osmocote Exact Hi-end, Scotts Company LLC). For each maternal plant or gene bank accession replicate, we have sown seeds into 3 plugs (one plug for each treatment), each plug with 3 seeds, to ensure that we would obtain at least one individual in each plug. We placed the seeded trays in a vernalization chamber at a constant temperature of 4°C from October until March and watered as required. The plants received, on average, 7 hours of light per day. When the seeds had germinated, we thinned the seedlings to retain a single individual per pot. In March 2024, we transferred the seedlings from the vernalization chamber to open foil tunnels in Düsseldorf and Marburg for several days to acclimate them to the outdoor conditions. We then transplanted one plant per genotype into each treatment: Düsseldorf soil, Marburg soil, and Marburg sand.

At each experimental site, we arranged the pots in a randomized incomplete block design to ensure that one genotype per population and, in Marburg, both soil treatments were represented within each block. We watered the plants daily using a drip irrigation system, except on days with rainfall.

We determined seven plant functional and life history traits. Flowering time, reproductive investment, and seed weight represent reproductive traits, while plant height, flag leaf area, specific leaf area, and total biomass represent growth-related traits. Additionally, we recorded reproductive biomass as a proxy of fitness. We recorded flowering time for each individual as the day of the year (DOY) of first spike emergence, defined as the stage at which the awns became visible through the base of the flag leaf (“heading”). Post-anthesis, we measured the height as the length of the longest tiller of each plant, excluding the spike. We then randomly selected three tillers with fully expanded spikes, harvested their flag leaves, and scanned the fresh leaves (EPSON Perfection V39). We then used the *pliman* package to determine the flag leaf area (Olivoto, 2022). We subsequently dried the leaves, weighed them, and calculated the specific leaf area (SLA) as total leaf area divided by total leaf dry mass.

After the trait measurements, we bagged the plants in air-permeable microperforated plastic bags to avoid seed loss during ripening. In June, when the majority of plants reached senescence, we harvested all plants and dried them at 40°C for 4 days. We subsequently separated the vegetative (tillers with leaves) and reproductive (spikes) biomass and weighed them, and calculated the reproductive investment as reproductive biomass divided by total biomass. In Düsseldorf, seed weight was determined by sampling 30 seeds per plant after harvest and scaling their seed mass to thousand-seed weight. In Marburg, seed weight was determined as thousand-seed weight by weighing and counting seeds obtained from 2 spikes that were previously covered with glassine bags to prevent cross-pollination.

### Statistical analysis

First, to understand the relationship among all traits, we conducted Pearson correlation analyses separately for each treatment. We visualized significant correlations using correlation matrices generated with the *Hmisc* package (Harrell Jr, 2025) (Figure S2). All correlation analyses were corrected for multiple testing using Benjamini-Hochberg correction (α = 0.05) (Benjamini & Hochberg, 1995).

Second, we tested the degree to which the variation in individual traits is heritable (that is, determined by maternal plant, population identity, or interaction between populations and treatment) or driven by environmental variation to treatment. To do this, we related reproductive (flowering time, seed weight, reproductive investment) and growth-related traits (plant height, flag leaf area, specific leaf area, total biomass) to treatment as a fixed factor and population identity, maternal plant identity nested within population, and the interaction between populations and treatment as random factors in linear mixed-effects models using the *lmer* function from *lme4* package (Bates et al., 2015), in total seven models, each for one response variable. We modeled the interaction between populations and treatment as a random grouping factor rather than a random slope because our goal was to partition variance into discrete components (population, maternal plant, interaction between populations and treatment) rather than estimate population-specific reaction norms. To estimate the proportion of variance explained by the fixed (treatment) and random (population, maternal plants nested in population, interaction between population and treatment) factors, we used the *r2()* function from the *performance* package (Zhang, 2016). To extract variance components for the random effect, that is, variance explained separately by populations, maternal plants nested within populations, and populations and treatment interaction, we used the *VarCorr()* function from the *lme4* package (Table S2). For this analysis, we excluded the gene bank accessions because no population-level replications were available.

Third, because we detected heritable variation in all traits, we tested which environmental factors explain this heritable trait variation. To do so, we isolated variation independent of experimental treatment by regressing trait values against treatment and used residuals from these models as response variables in the subsequent analyses. We then built seven linear mixed-effects models (one for each trait), using the residuals of each trait as the response variable, which we related to the environmental conditions at the seed collection sites. Specifically, we used two climatic predictors (mean annual temperature and annual precipitation) and three local environmental variables (vegetation cover, soil pH, and bioavailable soil phosphate) as fixed factors. In exploratory analyses, we found a quadratic relationship between temperature and some traits (Table S3). We thus added a squared temperature term in these models and retained the quadratic term when the model (ANOVA) comparison indicated a significantly better fit (Table S4). Population identity and maternal plant identity nested within a population were included as random effects to account for the non-independence among plants from the same population and the same mother plant. To determine the proportion of variation explained by the model terms, we used the *r2()* function from the *performance* package. The relative importance of individual fixed-effects predictors was assessed using the *partR2()* function from the *partR2* package (Stoffel et al., 2021). Note, because interactions between populations and treatment explained only a small proportion of trait variation, we focused these analyses on heritable differences in mean trait values rather than heritable plasticity (G×E). We did not use gene bank accessions for these analyses because there were no environmental variables available for the collection sites (except climate). Because mean annual temperature at the seed collection site was the main predictor of heritable trait variation, we visualized this relationship (Figure S3).

We further explored whether the relationships between plant traits and mean annual temperature at the seed collection sites were consistent between the measurements made in the common gardens and those made *in situ*. To do this, we related *in situ* traits (date of seed ripening as a proxy of reproductive phenology, plant height, and seed weight) to mean annual temperature in a linear mixed-effects model for each trait, using the *lme4* package, and visualized them alongside the data for corresponding traits from the common garden. Note that a direct comparison in one model was not possible because the traits were measured slightly differently *in situ* and in the common garden. For example, phenology in the common garden was recorded on an individual level as the onset of flowering, but *in situ* as the time of seed ripening on the population level.

Fourth, we tested whether variation in fitness reflects broad-scale temperature adaptation. To do this, we tested whether plants achieved highest fitness when their climate of origin closely matches the growing conditions of the common gardens. Note, because the plants in both common gardens experienced growing conditions from March to June, we used the average spring temperature from March to June for the years 2000–2020. We used reproductive biomass as a proxy for individual fitness and examined its relationship with the mean spring temperature (SpringT) of the population’s origin for each treatment. Because this relationship followed a unimodal pattern, we modeled the fitness-temperature relationships using a quadratic mixed-effects model. To test whether the temperature of origin associated with maximum fitness differed among the soil treatments and the common garden sites, we fitted the quadratic mixed-effects model including an interaction term:

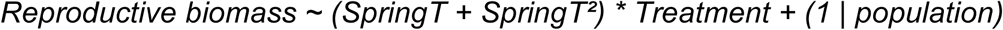

For each treatment, we estimated the temperature of origin associated with maximum fitness (i.e., the SpringT at seed origin that would produce maximal reproductive biomass in a given treatment) as the peak of the fitted quadratic curve. We defined the temperature offset as the difference between the optimum origin-temperature and the mean spring temperature recorded at the experimental site (optimum origin-SpringT minus garden SpringT), with positive values indicating that maximum fitness was associated with populations originating from warmer climates than the garden itself. To evaluate the robustness of the estimated optima, we calculated confidence intervals by using two resampling approaches. First, we performed parametric bootstrapping of the fitted mixed-effects model using the function *bootMer* (1,000 bootstrap samples) from *lme4* package. The model was specified as:

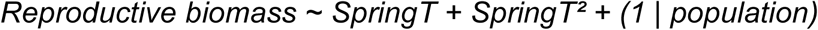

where the mean spring temperature (SpringT) of the plants’ origin was included as a fixed effect with a quadratic term, and population identity was included as a random intercept. We then conducted model-based simulations using the *simulate* function (200 repetitions) from base R, generating response values based on the fitted model, using the SpringT values, and population-level random effects drawn from the estimated variance components. These complementary resampling approaches consistently identified similar peak values across all treatments (Table S5). We then compared the estimated fitness optima from all three treatments with SpringT to assess whether peak fitness occurred under home-temperature conditions, as inferred from the mean spring temperature between 2000 and 2020.

To assess model assumptions, we evaluated the model fit using residual diagnostics (Zuur et al., 2010). Response variables were log-transformed where necessary. We conducted all data analyses using R version 4.5.3 (R Core Team, 2026) within the RStudio environment.

## Results

Flowering time and plant height were positively correlated in all three treatments (Figure S2). In contrast, the relationship between flowering time and fitness, approximated by reproductive biomass, varied among treatments: Flowering time was positively correlated with fitness in the Marburg soil treatment (r = 0.15), negatively correlated in the Düsseldorf soil treatment (r = -0.15), and showed no significant correlation in the Marburg sand treatment.

Variation of all traits was significantly affected by treatment (E, phenotypic plasticity), identity of maternal plant and population (G, genetic differentiation), and in some traits also by population and treatment interaction (GxE, heritable plasticity). Although the proportion of variance explained by each factor differed between traits, we did not observe any consistent differences between reproductive and growth-related traits. Flowering time showed the highest heritable contribution, followed by plant height, while flag leaf area and total biomass were mostly influenced by the treatments (plastic responses). Heritable trait variation was primarily associated with population identity (4-68 %), while maternal plant identity contributed smaller but consistent proportions, with variation explained by 1.3-9 %, except for total biomass. The interaction between population and treatment (GxE) contributed to five out of seven traits, with 0.9-9 % variation explained, which is less than population and maternal plant identity (G) (Figure 2, Table S2).

**Figure 2.**
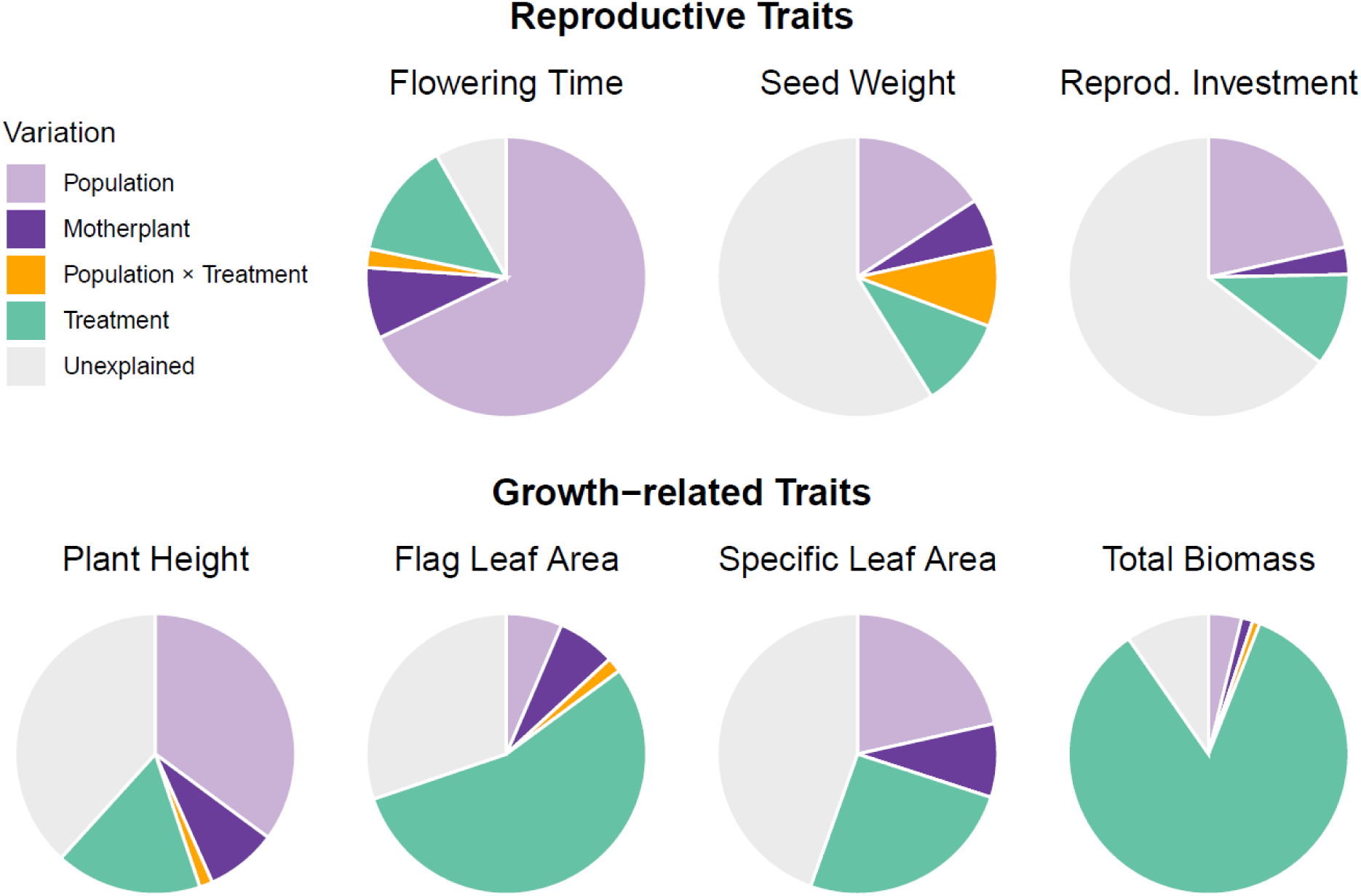
Partitioning of trait variation measured in the common garden experiments as explained by population, mother plant nested within a population, interaction between population X treatment, and treatment (see Table S3). Gene bank accessions were not included.

The environment at the site of seed origin explained 0-48 % of residual trait variability in the common garden experiment, after correcting for the experimental treatment (Figure 3). The main driver was mean annual temperature, which explained 3-44 % of variation in 5 out of 7 traits, specifically in flowering time (44 %), plant height (27 %), SLA (16 %), reproductive investment (14 %), and total biomass (3 %). Mean annual precipitation did not explain a significant proportion of variation in any of the traits. Local vegetation cover and soil conditions at seed collection sites contributed comparatively little to trait variation (Figure 3, Table S2). In all trait-environment models, population and maternal plant identity still explained a substantial proportion of variation, indicating that trait variation is strongly affected by genetic differentiation even after correcting for *in situ* environmental variation (Figure 3, Table S2)

**Figure 3.**
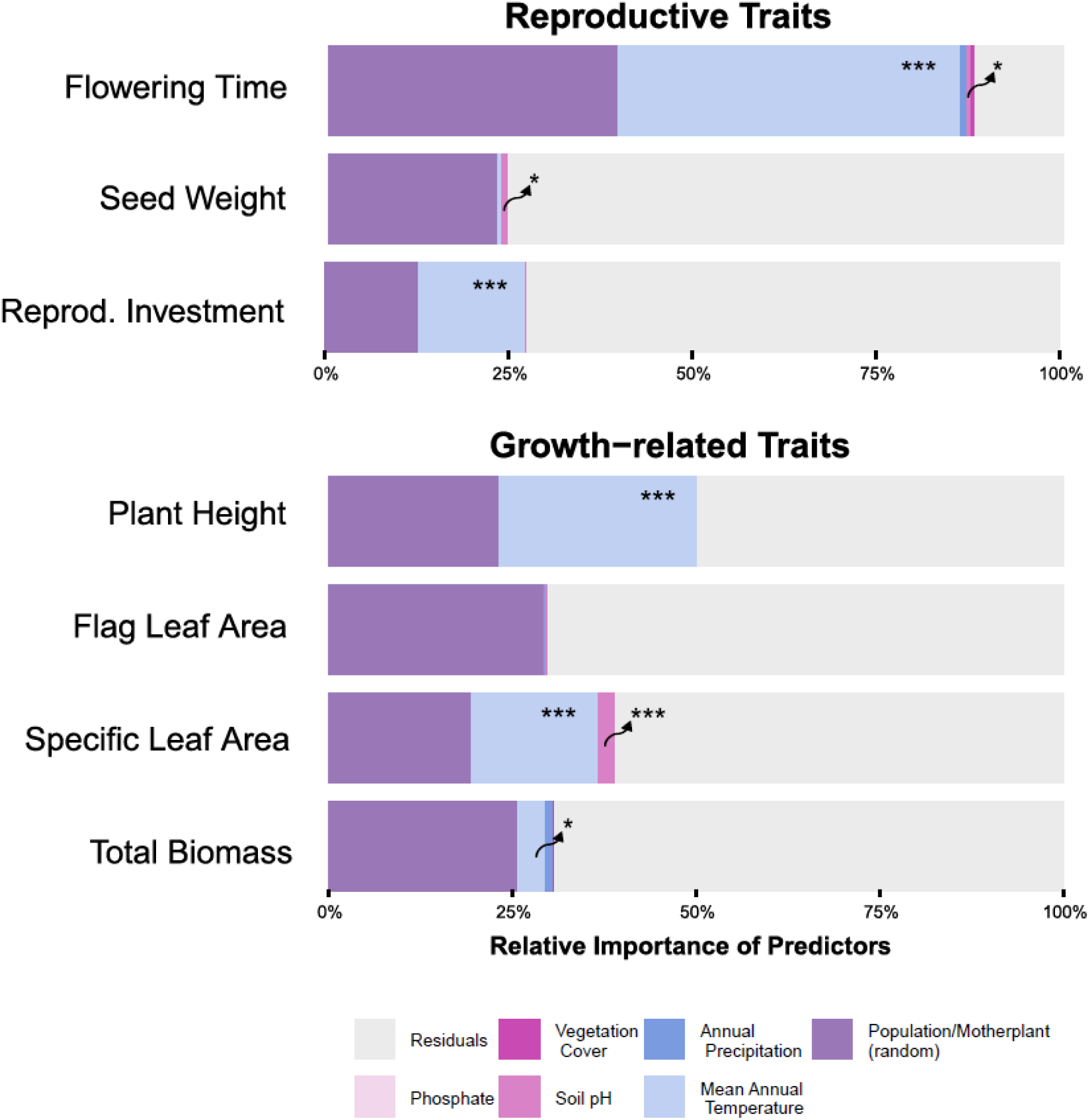
Trait variation in the common gardens explained by environmental conditions in the seed origin: climate (mean annual temperature and annual precipitation), competition (vegetation coverage), and soil (pH-value and bioavailable phosphate) as fixed factors, and population and mother plant identity nested within a population as random factors. The treatment effects were accounted for before estimating the model. Gene bank accessions are not included.

In the common garden, plants originating from warmer regions flowered earlier and were smaller (Figure 4). They also produced lower biomass, had lower SLA, and higher reproductive investments, although these relationships were quadratic. Seed weight showed a weak quadratic relationship, with a peak in plants from areas with a mean annual temperature around 16°C. The observed patterns were largely consistent across treatments (Figure S4). The trait variation in the common garden along mean annual temperature partly reflected patterns observed *in situ*, particularly for reproductive phenology and plant height, but not for seed weight (Figure 4, Table S4).

**Figure 4.**
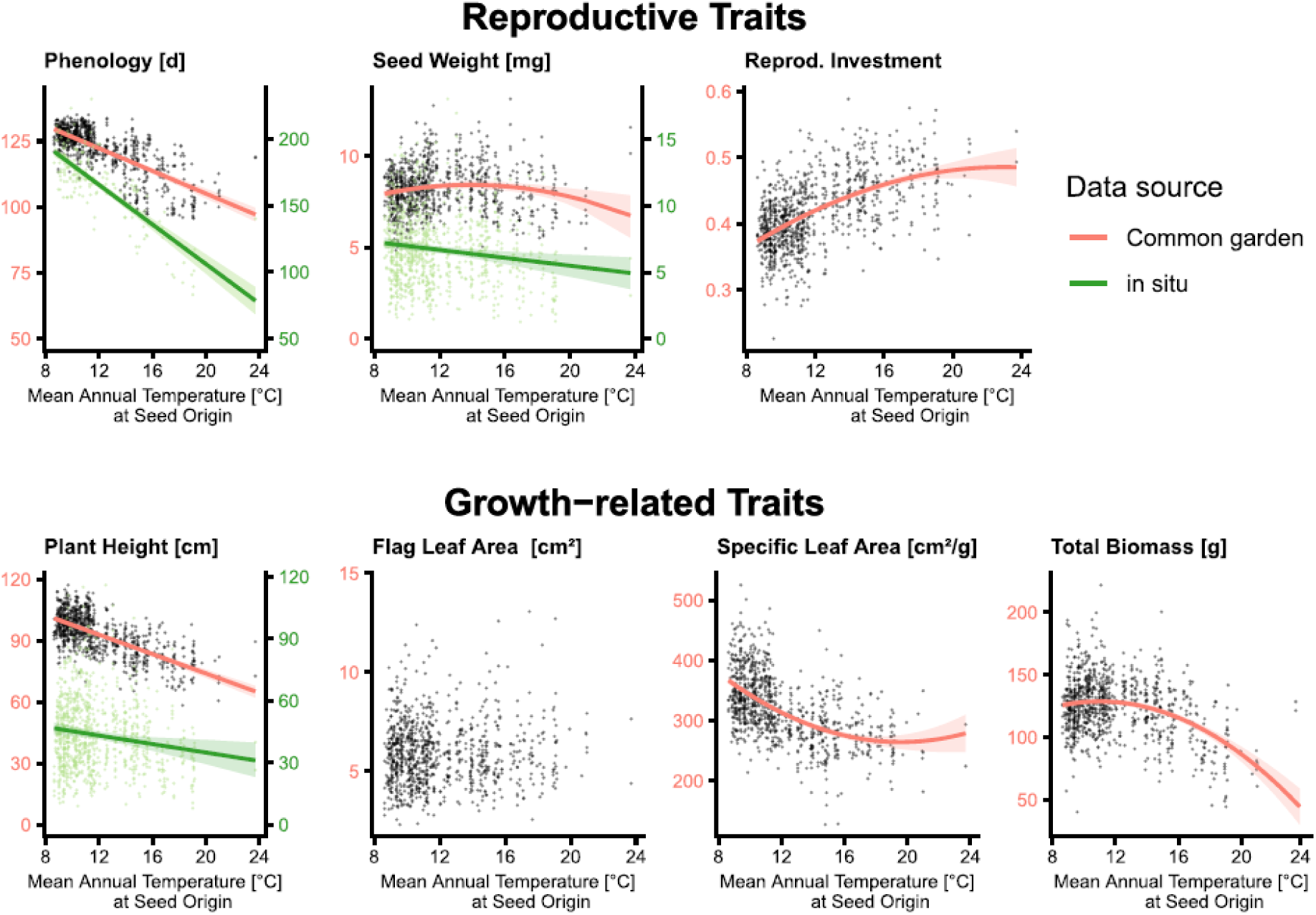
Relationship between trait variation and temperature in the plants’ origin. Phenology (flowering time in the common garden, seed ripening onset in situ), seed weight, and plant height were sampled *in situ* (green regression line) and in the common gardens (red regression line). Common garden traits are averaged by maternal plant identity across treatments because the trait patterns were largely consistent among treatments (Figure S4).

Plants did not achieve the highest fitness, as approximated by reproductive biomass, under temperatures similar to their own site of origin (Figure 5 A-C). The detected significant interactions between treatment and the quadratic temperature term indicate that the estimated optimum-origin temperature differed among experimental environments (Table 1). Under fertilized soil conditions in both Marburg and Düsseldorf, the estimated optimum corresponded to populations originating from regions approximately 2°C warmer than the mean spring temperature at the experimental sites (Figure 5, Table S5). In the sand-based treatment, the temperature offset was even greater, with the estimated optimum corresponding to populations from regions approximately 4°C warmer. (Table S5, Figure 5).

**Figure 5.**
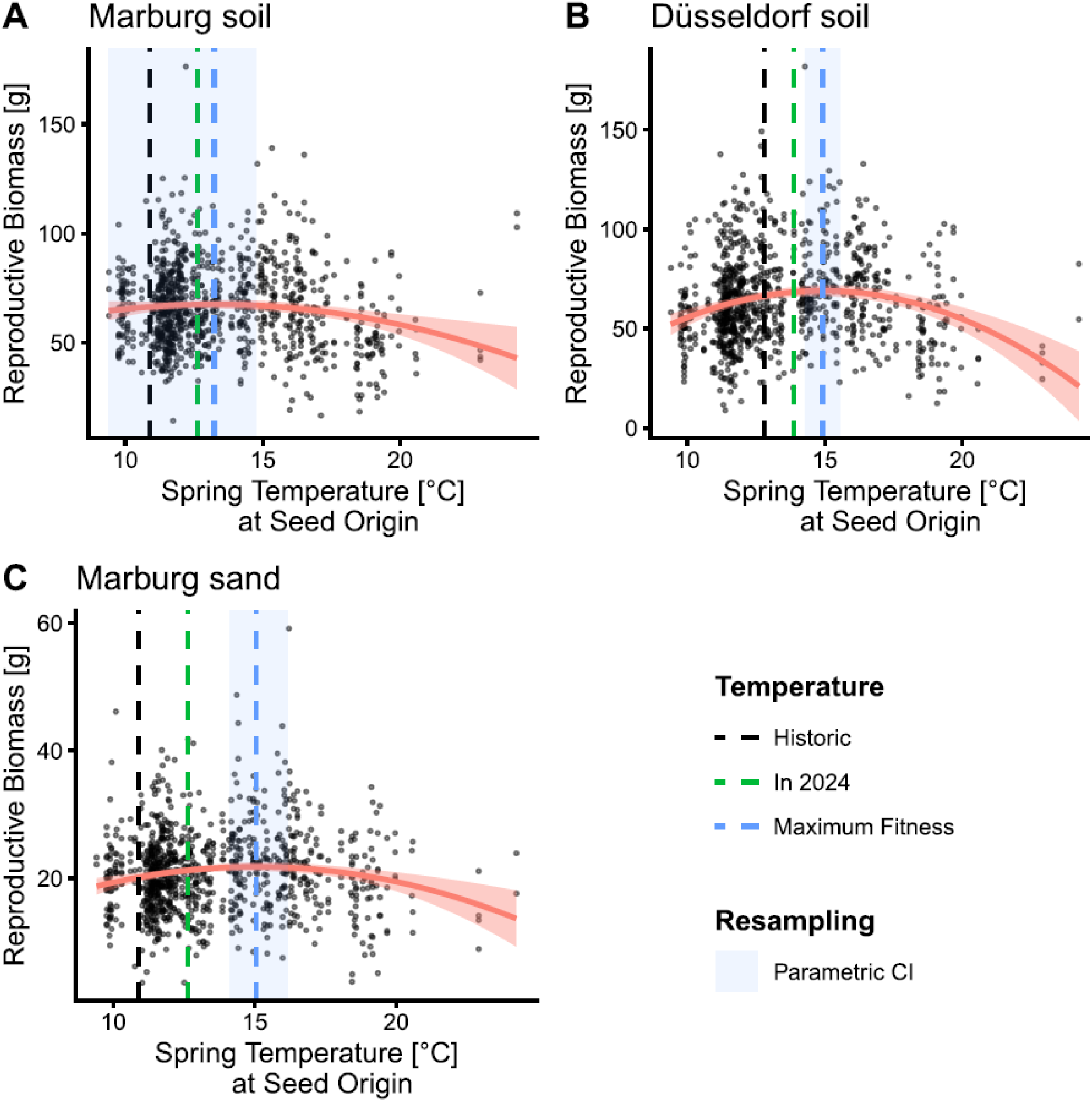
Relationship between the fitness proxy (reproductive biomass) and mean spring temperature at the population’s site of origin. Blue dashed lines indicate the estimated fitness optimum (peak of reproductive biomass), green dashed lines display the estimated spring temperature in 2024, and black dashed lines represent the historic mean spring temperature at each common garden site. Blue-shaded areas denote confidence intervals derived from parametric bootstrapping, used to assess the robustness of the estimated optima.

**Table 1.** The relationship between fitness (reproductive biomass) and mean spring temperature (Spring Temp). Results of a Type III analysis of variance (ANOVA) for a linear mixed-effects model, including a quadratic temperature term and its interaction with treatment. Statistically significant effects are shown in bold (p < 0.05).

| Reproductive Biomass | df | Df.res | F | *p* |
| --- | --- | --- | --- | --- |
| Spring Temp | 1 | 498.64 | 25.98 | **< 0.001** |
| Spring Temp<sup>2</sup> | 1 | 626.81 | 55.45 | **< 0.001** |
| Treatment | 2 | 2732.18 | 1781.00 | **< 0.001** |
| Spring Temp:Treatment | 2 | 2733.23 | 15.86 | **< 0.001** |
| Spring Temp<sup>2</sup>:Treatment | 2 | 2732.51 | 21.77 | **< 0.001** |

## Discussion

To disentangle heritable variation from phenotypic plasticity and assess adaptive trait variation and potential lags under climate change, we combined *in situ* data with common garden data. Contrary to our initial expectations, we found that heritable variation (estimated as the proportion of trait variance attributed to population identity, maternal plant identity, and populations and treatment interaction) was not restricted to reproductive traits but was also pronounced in growth-related traits, indicating that heritable contribution may instead relate to the degree of genetic control over each trait’s development than on its functional category. Flowering time and plant height showed substantial heritable components, whereas seed weight, reproductive investment, and SLA had intermediate heritable contributions. In contrast, flag leaf area and total biomass were primarily driven by plastic responses to the experimental treatments.

Heritable variation was mainly associated with temperature at the populations’ origin, which is consistent with patterns observed *in situ*. Plants from warmer regions flowered earlier, reached the onset of seed ripening sooner, and remained smaller, indicating strong directional selection along thermal gradients. However, our results also suggest that adaptation may lag behind ongoing climate change.

### Heritable differentiation and phenotypic plasticity

All measured traits were shaped by both heritable variation and plastic responses, although to different degrees. We expected reproductive traits to be predominantly determined by differentiation between maternal plants and populations and thus be heritable, and growth-related traits to be mainly driven by phenotypic plasticity (Villellas et al., 2021). However, our results indicate that heritable contribution to trait variation does not strictly follow this distinction, which contrasts with previous study on a short-lived plant *Plantago lanceolata* (Villellas et al., 2021).

The heritability of a trait is possibly more linked to its developmental regulation than to the ecological function (Falconer & Mackay, 1989). Flowering time, for example, is under strong genetic control through key regulatory genes (Hill & Li, 2016), while plant height, though similarly heritable, reflects the cumulative action of many loci of small effects (Mikołajczak et al., 2017). The strong correlation we observed between flowering time and plant height likely reflects developmental dependency because plants that flower later have more time to grow taller. In barley, some flowering time genes can exert pleiotropic effects (one gene affects several traits) on plant height, which may further contribute to this correlation (Teplyakova et al., 2017). In contrast, total biomass integrates resource acquisition over the entire growing season, while flag leaf area reflects resource capture at a specific developmental stage, and both are more responsive to local environmental conditions (Matesanz et al., 2020), which likely explains their lower heritability. Seed weight, reproductive investment, and SLA showed intermediate heritable contributions, reflecting a combination of genetically determined allocation strategies and environmentally sensitive resource availability during their development (Scheepens et al., 2010); for instance, seed provisioning depends on genetically influenced maternal allocation patterns but is ultimately constrained by resources accumulated under the prevailing treatment conditions (Lorts et al., 2020). Together, our results suggest that the degree of genetic determination of a trait is better predicted by its evolutionary history and environmental sensitivity than by a simple distinction between reproductive and growth-related traits.

### Drivers of heritable trait variation

Heritable trait variation in *H. murinum* was driven primarily by broad-scale climatic gradient, specifically by mean annual temperature at seed origin but not precipitation. The driving effect of mean annual temperature on heritable trait variations is consistent with previous studies (Griffin-Nolan et al., 2025; Macel et al., 2007). However, the lack of precipitation effects on heritable differentiation is surprising because many studies found precipitation to be the driving force (Britton et al., 2026; Kilkenny et al., 2026; Møller et al., 2026), sometimes exceeding the effect of temperature (Oyarzabal et al., 2008). A possible explanation is that the relationship between precipitation and plant-available water is indirect, as water availability is shaped by a combination of factors, including rainfall seasonality, hydrology, soil depth, and type, and this is also partially mediated by temperature (Moles et al., 2014).

Plants from warmer regions were smaller, produced less biomass, flowered earlier, and invested more in reproduction. We observed a similar pattern *in situ* and in the common garden, which indicates that these trait syndromes are under genetic control and likely reflect adaptive variation by directional selection along climatic gradients (Yan et al., 2021). Plants from warmer areas flower earlier and thus have less time for vegetative growth, resulting in lower biomass and plant height compared to later-flowering conspecifics. Accordingly, in hot and dry regions, plants must prioritize reproduction over growth earlier in their life to reproduce before drought and heat-induced mortality (Nour et al., 2024), especially in annual plant species. In contrast, plants from colder, more humid regions can delay flowering and extend the vegetative growth period, thereby maximizing fitness through greater height and biomass. This differential selection for flowering time was reflected in the two common gardens, where flowering time was positively correlated with fitness in Marburg in ambient soil, but negatively correlated in the 2°C warmer climate of Düsseldorf. A similar heat escape strategy through advanced flowering has been documented in other early-flowering species (Ehrlén et al., 2023; Moore & Lauenroth, 2017). The decrease of SLA towards warmer regions also suggests adaptation to heat and drought through increased water-use efficiency (Petrík et al., 2024) and might also indicate enhanced leaf thermoregulation, as leaf structural traits can help maintain leaf temperature within an optimal range under heat stress (Michaletz et al., 2015).

In contrast to climate, local environmental conditions like soil properties and competition had only negligible effects on heritable trait variation. One possible explanation is that broad-scale climate represents a spatially structured and temporally consistent selective pressure, whereas local soil and biotic conditions vary more finely (Briscoe Runquist et al., 2020). Given *H. murinum*’s effective long-distance dispersal, gene flow among populations may outpace this fine-scale variation, preventing effective local adaptation to smaller spatial scales (Bontrager et al., 2020). The strongest effect we detected was a decrease of SLA with soil pH at the seed origin, which explained 2.3 % of the heritable variation in this trait. However, plant adaptation to pH is not common across species (Rupprecht et al., 2021), likely because pH itself is not a direct selective agent but rather a proxy for a range of downstream chemical conditions it governs.

### Plant performance under climate change

We found that fitness, approximated by reproductive biomass, peaked in populations with a temperature offset of 2-4 °C, i.e., originating from climates warmer than the mean spring temperature in the respective common gardens. This suggests that current temperature adaptation of the populations is not keeping pace with ongoing environmental changes. In fact, plants faced elevated temperatures during the experiment because the mean spring temperature at our common garden sites was 1.1 and 1.7 °C, respectively, warmer during the experimental year (2024) than the historical baseline used to define climatic adaptation (mean of 2000-2020, Table S5), which may partly explain the advantage of warm-origin populations (Wilczek et al., 2014). This temperature offset is comparable in magnitude to lags reported in other species, including *Arabidopsis thaliana* and *Trifolium repens* (Albano et al., 2026; Wilczek et al., 2014). Originally, we hypothesized that *Hordeum murinum* would track climate change effectively because of its high dispersal potential (Krause et al., 2015). However, the observed mismatch between the temperature of origin associated with maximum fitness and contemporary spring temperatures indicates that dispersal alone has not been sufficient to fully track ongoing climatic change, a pattern that corresponds to broader evidence (Román-Palacios & Wiens, 2020).

The advantage of warm-adapted genotypes could be partially enhanced because we grew plants in pots. In pots, roots experience stronger temperature fluctuations than when plants grow directly in the soil and thus face higher temperatures in spring and summer. On the other hand, we have watered the plants, which reduced the hydric stress that is connected to elevated temperatures. Future studies incorporating natural soil and field conditions are therefore needed to verify the extent to which these patterns reflect adaptation *in situ*.

Plant performance could theoretically be affected by maternal effects because we worked with wild-collected plants without a refresher generation. This might have contributed to the variation explained by the maternal plant. However, the variation was small compared to that explained by population identity. Additionally, maternal effects are genotype-specific and likely to introduce noise, rather than be behind the large-scale pattern we detected (Latzel et al., 2023).

## Conclusions

Variation in plant functional and life history traits in *Hordeum murinum*, an annual ruderal grass, is shaped by both plasticity and heritable variation. This heritable variation is shaped predominantly by mean annual temperature, while local edaphic factors and plant-plant competition play only a minor role. However, our results indicate that dispersal-mediated climate tracking may not keep pace with rapid environmental shifts, leaving populations climatically mismatched with current conditions at their site of origin. Future studies are needed to incorporate natural environmental conditions to verify the magnitude of adaptation.

## Acknowledgements

We thank Shahal Abbo, Maximilian Hanusch, David Lafleuriel, Pierre-Marie le Henaff, Richard Oliver, Soraya Safavi, Malte Schmidt, and Lena Ulber for collecting samples, and Lisa Grunwald, Liliane Irle, Jonas Lampe, Laura Libera, Christina Mengel, Freya Müller, Sarah Paschen, and Nadine Pluquette for technical support.

## Authors’ contribution

AB and MK conceived the idea, AB, MK, SJL, HV, TH, and SL designed the study. HV analyzed the data and wrote the first draft. All authors collected data, edited the manuscript, and approved the submission.

## Supplementary information

**Table S 1.**
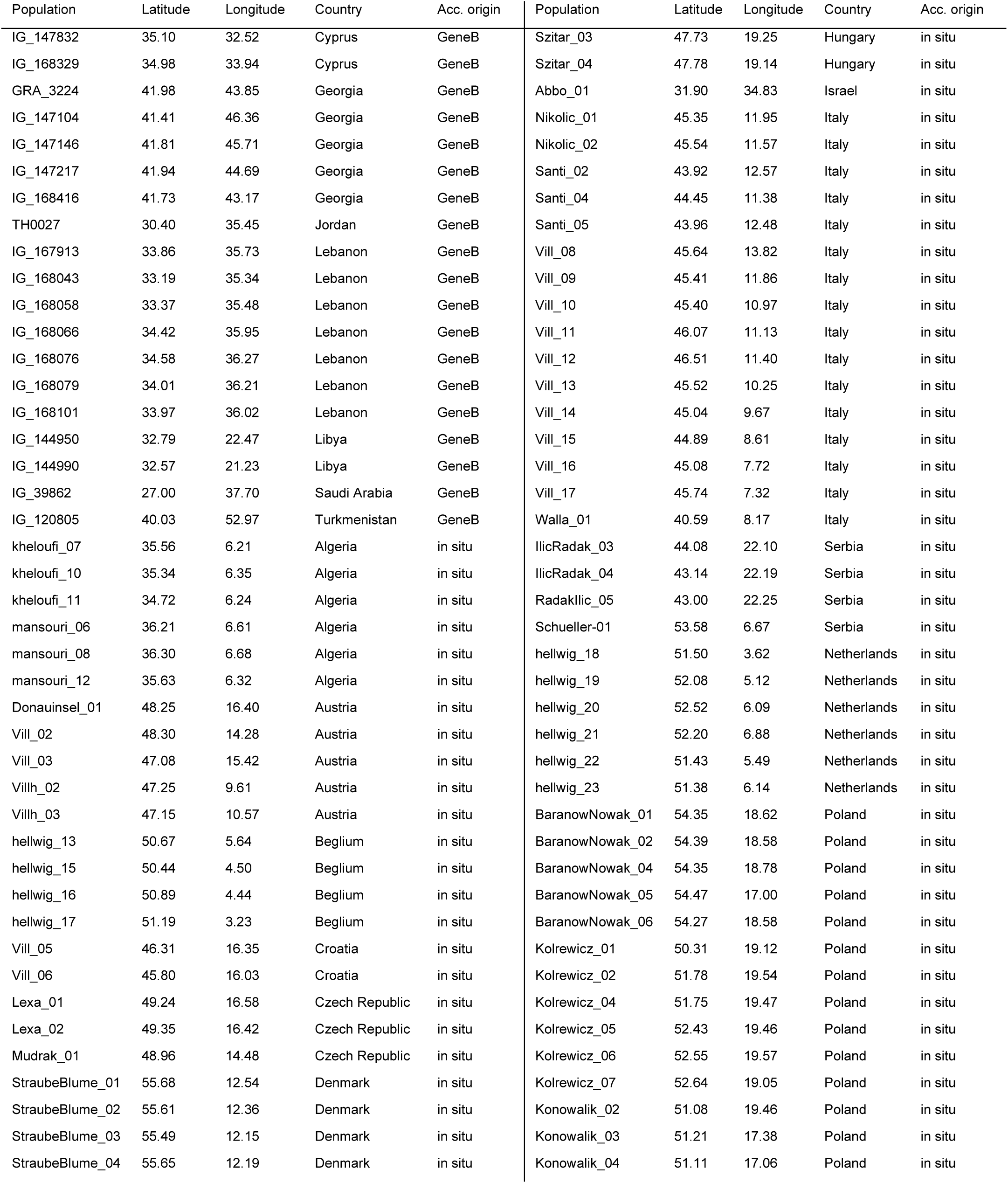
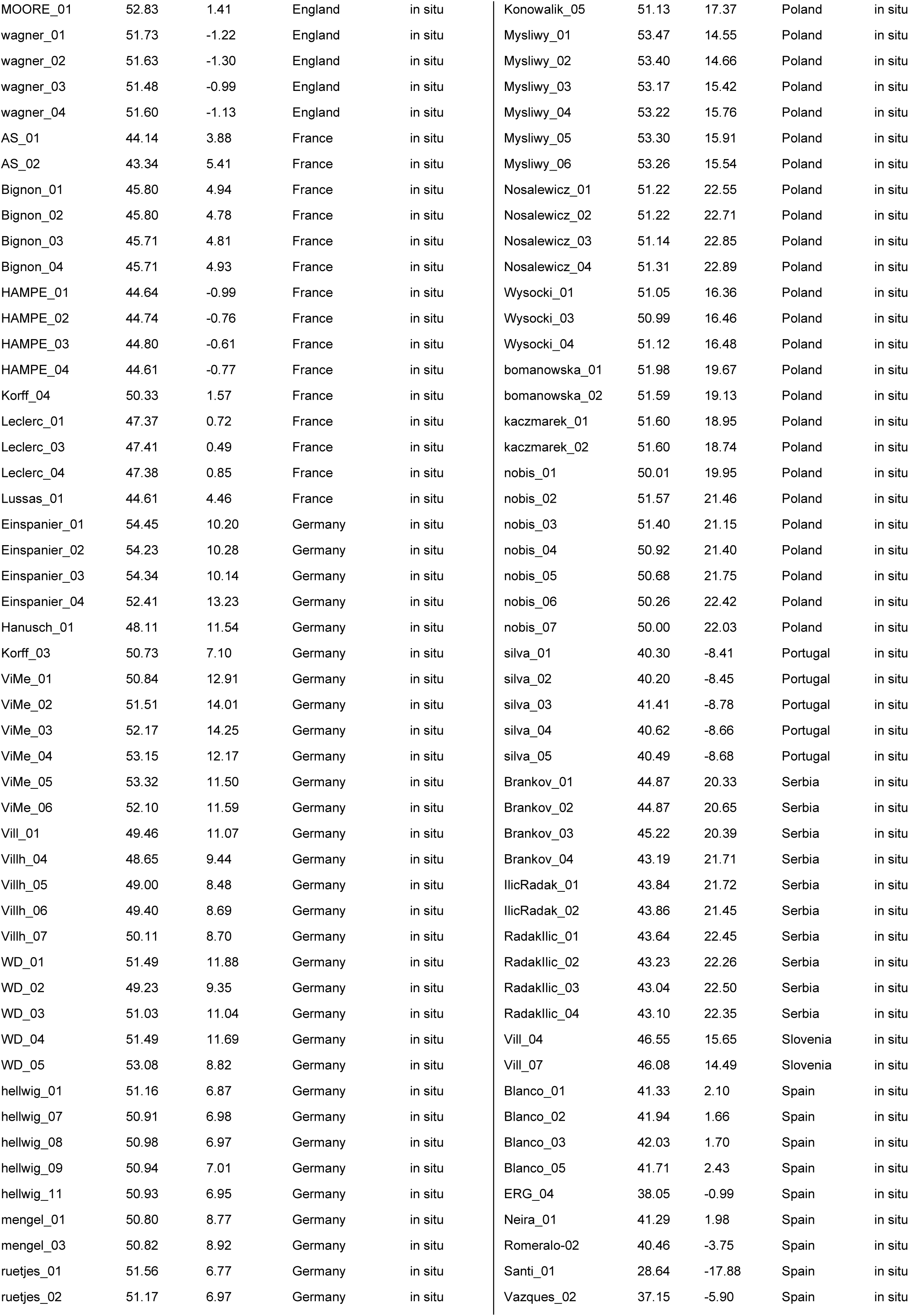
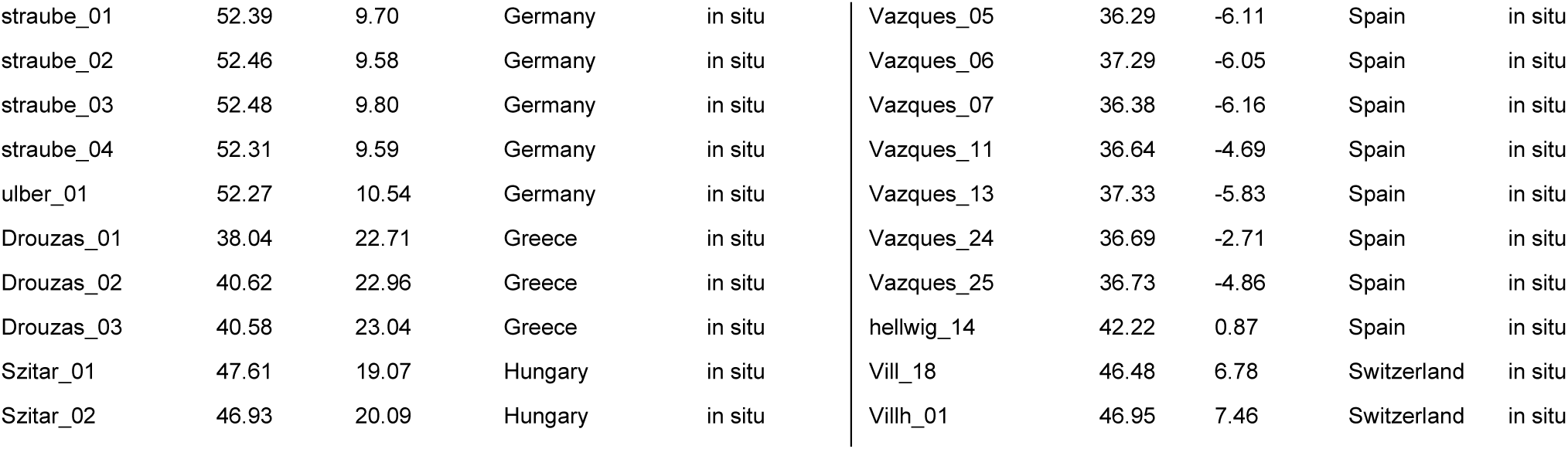
List of all populations used in the study: accession origin (Acc. Origin) from gene bank (GeneB) and populations sampled in the field (in situ). All coordinates (Latitude and Longitude) were averaged at the population level.

**Figure S 1.**
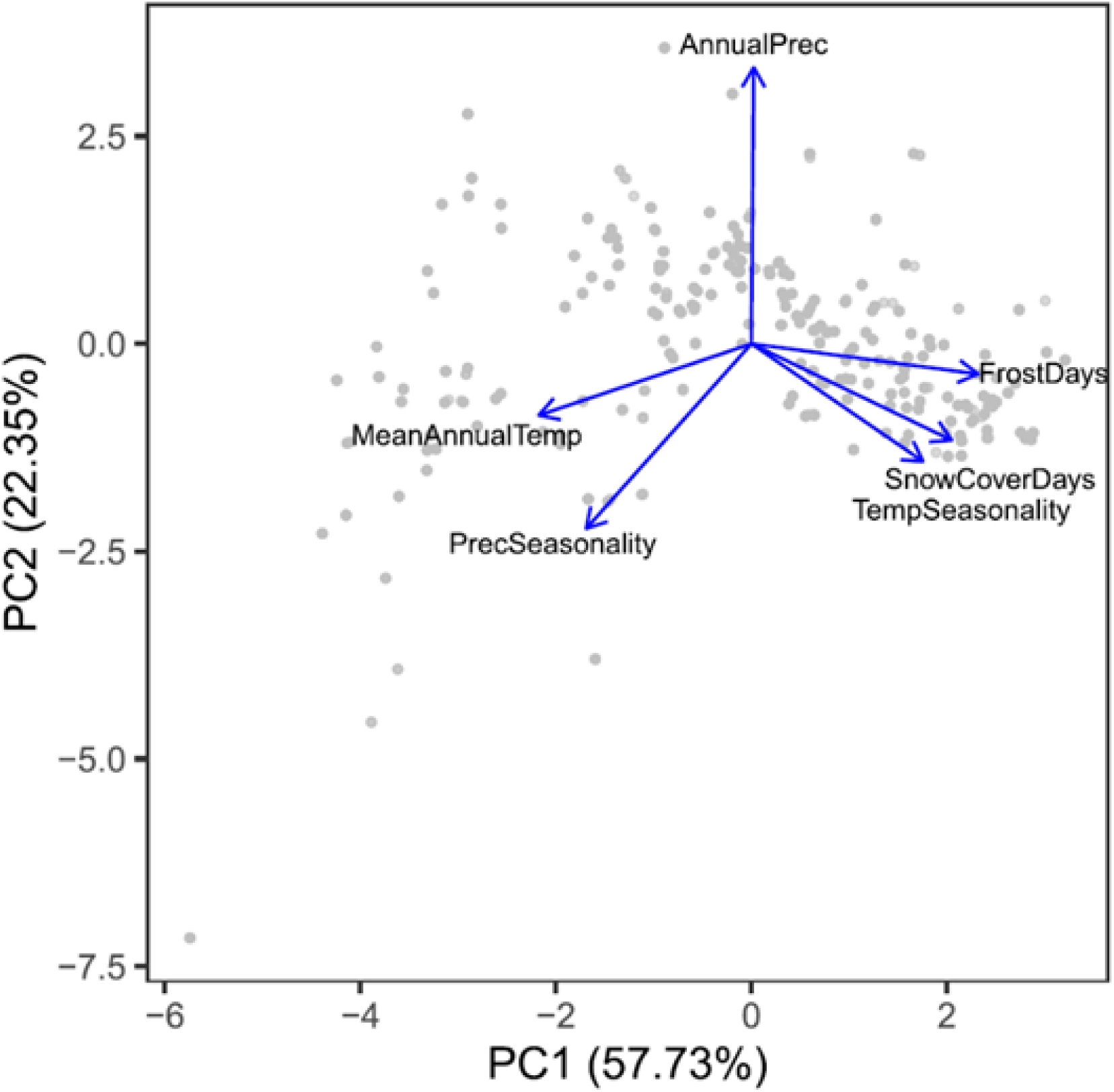
Principal component analysis (PCA) of the climatic variables derived from the CHELSA dataset, with PC1 explaining 57.73% and PC2 explaining 22.35% of the total variance. Climatic variables (blue arrows) indicate the direction and strength of their contribution to the principal components. Longer arrows denote stronger correlations with the respective axes.

**Figure S 2.**
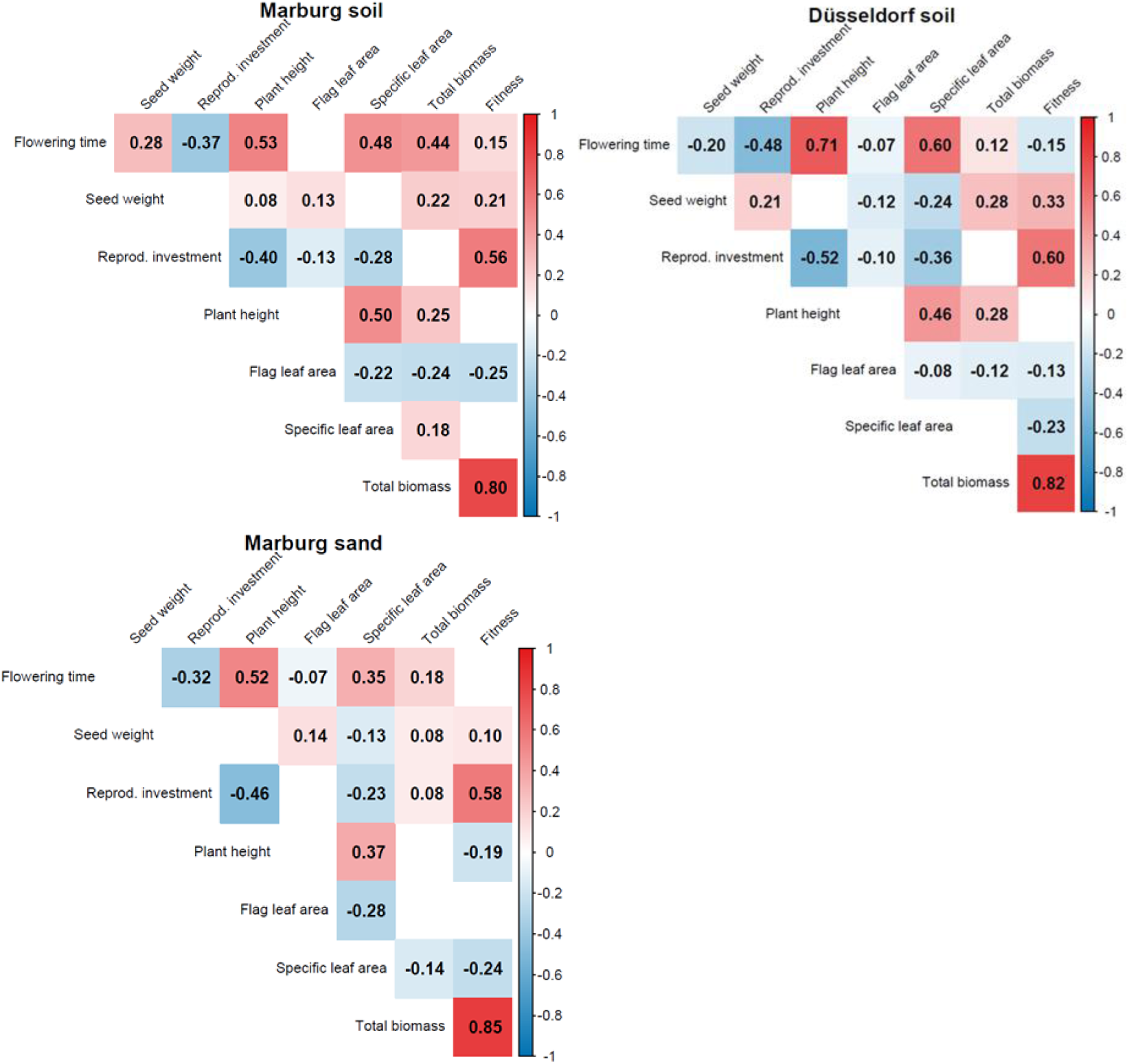
Pearson’s correlation coefficient between all studied plant traits separated by treatment (Düsseldorf soil, Marburg soil, Marburg sand treatment). The number and color intensity in the cells indicate the strength and direction of two related variables. Empty cells indicate non-significant correlations (p > 0.05). The significance threshold was adjusted for multiple testing using Benjamini-Hochberg correction.

**Figure S 3.**
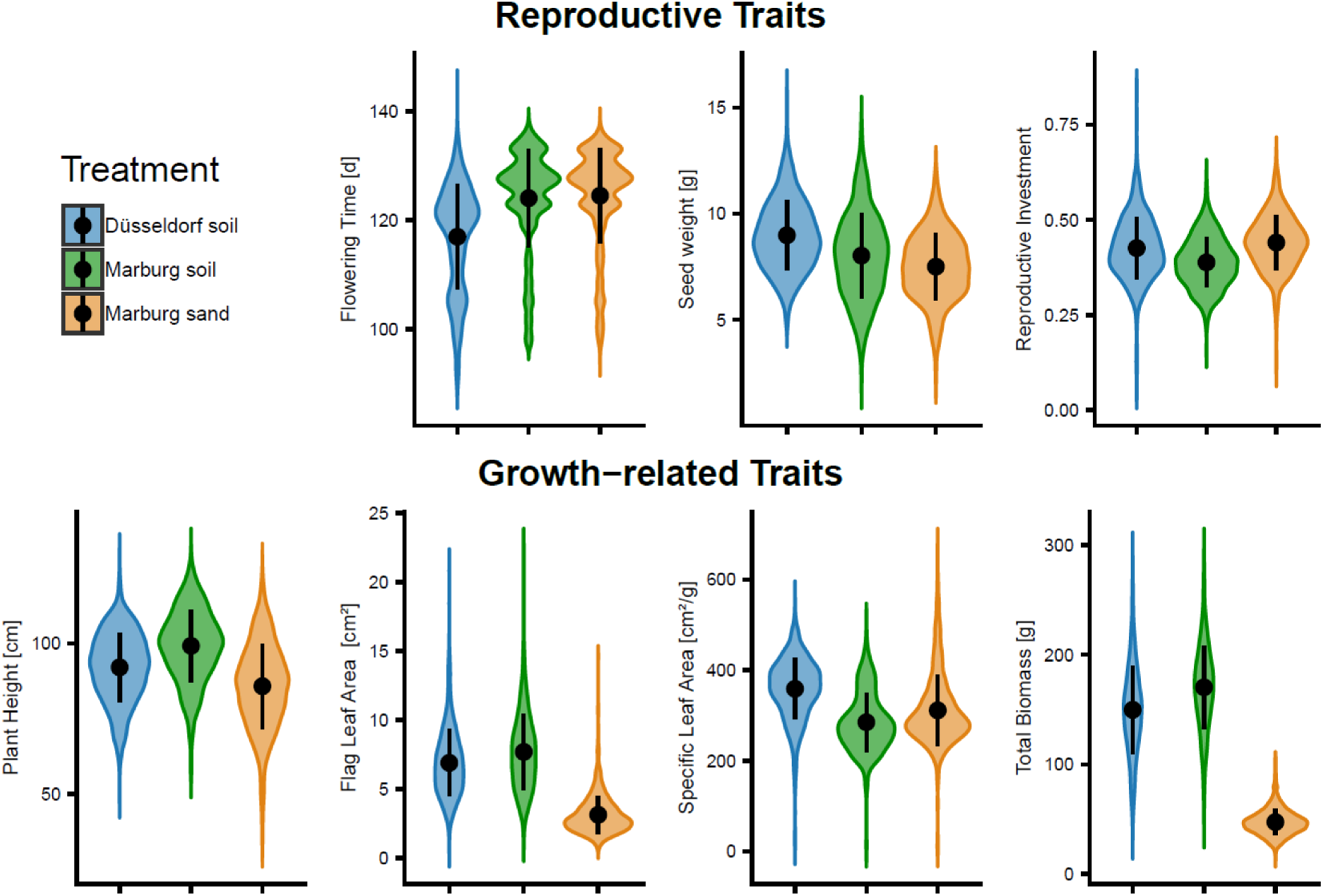
Trait variation across treatments for each measured trait. Violin plots illustrate differences among treatments (Düsseldorf soil in blue, Marburg soil in brown, Marburg sand in yellow) and the distribution of trait values within each treatment. The black dot and line represent the median (black dot) and the interquartile range (black line).

**Figure S 4.**
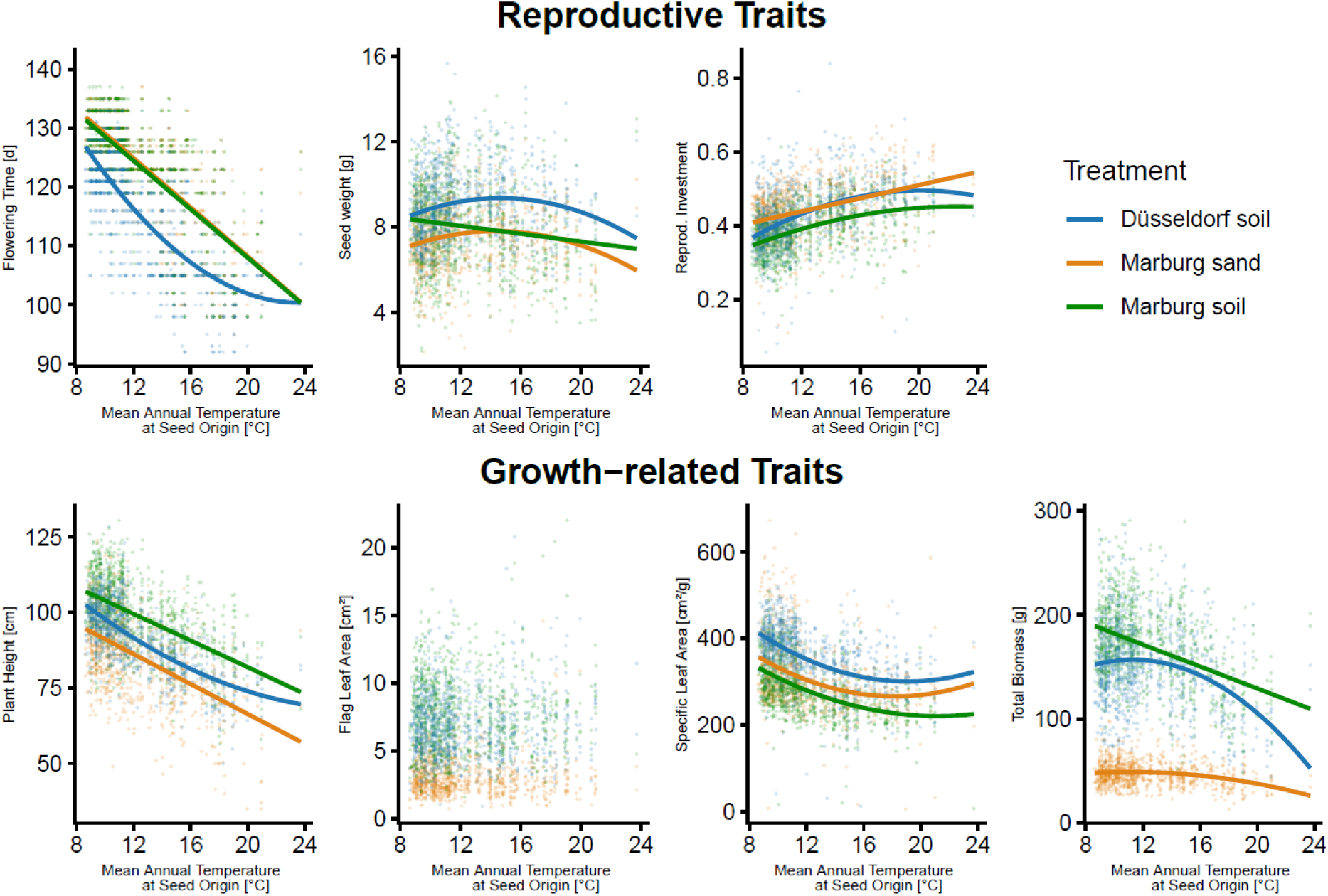
Linear or quadratic relationships between trait values and temperature of maternal plants’ origin separated by treatment (Düsseldorf soil: blue, Marburg soil: brown, Marburg sand: yellow). Regression lines are displayed only when relationships are statistically significant (p < 0.05).

**Table S 2.** Trait variation explained by treatment, population identity, and mother plants nested within populations. Results of linear mixed-effects models with traits as response variables, treatment as fixed effect, and population identity and maternal plants as random factors.

| Trait | Population | Motherplant | Population x |  | Unexplained | Condit. R<sup>2</sup> |
| --- | --- | --- | --- | --- | --- | --- |
|  |  |  | Treatment | Treatment |  |  |
| Flowering Time | 0.679 | 0.082 | 0.022 | 0.135 | 0.083 | 0.917 |
| Seed Weight | 0.158 | 0.057 | 0.092 | 0.104 | 0.589 | 0.411 |
| Reprod. Investment | 0.216 | 0.031 | 0.000 | 0.107 | 0.646 | 0.247 |
| Plant Height | 0.351 | 0.082 | 0.015 | 0.169 | 0.383 | 0.617 |
| Flag Leaf Area (log) | 0.064 | 0.067 | 0.017 | 0.549 | 0.302 | 0.698 |
| Specific Leaf Area | 0.215 | 0.085 | 0.000 | 0.254 | 0.446 | 0.300 |
| Total Biomass (log) | 0.038 | 0.013 | 0.009 | 0.844 | 0.097 | 0.903 |

**Table S 3.** Comparison of quadratic and linear relationships between trait variation and mean annual temperature. Results of linear mixed-effects models. Analysis of variance with error type II. Significant effects (p < 0.05) are highlighted in bold.

| Trait | Model choice p | Model | Estimate | Term | F | df | Df.res | p |
| --- | --- | --- | --- | --- | --- | --- | --- | --- |
| *Common garden* |  |  |  |  |  |  |  |  |
| Flowering Time [d] | **0.048** | quadratic | -3.93 | MAT | 18.54 | 1 | 220.31 | **<0.001** |
|  |  |  | 0.064 | MAT<sup>2</sup> | 3.88 | 1 | 223.95 | 0.05 |
| Seed Weight [mg] | **0.005** | quadratic | 0.48 | MAT | 8.05 | 1 | 232.14 | **0.005** |
|  |  |  | -0.02 | MAT<sup>2</sup> | 8.15 | 1 | 241.35 | **0.005** |
| Reprod. Investment | **0.004** | quadratic | 0.025 | MAT | 23.03 | 1 | 242.94 | **<0.001** |
|  |  |  | -0.001 | MAT<sup>2</sup> | 8.20 | 1 | 255.96 | **0.005** |
| Plant Height [cm] | 0.087 | linear | -2.37 | MAT | 383.73 | 1 | 227.55 | **<0.001** |
| Flag Leaf Area [cm<sup>2</sup>] | 0.625 | linear | 0.02 | MAT | 0.78 | 1 | 229.92 | 0.377 |
| Specific Leaf Area [cm<sup>2</sup>/g] | **<0.001** | quadratic | -33.52 | MAT | 36.11 | 1 | 245.94 | **<0.001** |
|  |  |  | 0.86 | MAT<sup>2</sup> | 18.17 | 1 | 259.88 | **<0.001** |
| Total Biomass [g] | **<0.001** | quadratic | 7.74 | MAT | 6.99 | 1 | 234.45 | **0.009** |
|  |  |  | -0.39 | MAT<sup>2</sup> | 13.52 | 1 | 244.51 | **<0.001** |
| *In situ* |  |  |  |  |  |  |  |  |
| Seed ripening [d]<br>(population-level) | 0.7645 | linear | -7.4367 | MAT | 79831<br>(sum sq) | 1 | 307.75 | **<0.001** |
| Seed weight [mg] | 0.1415 | linear | -0.15 | MAT | 8.75 | 1 | 191.47 | **0.004** |
| Plant height [cm] | 0.05 | linear | -1.019 | MAT | 8.82 | 1 | 191.06 | **<0.001** |

**Table S 4.**
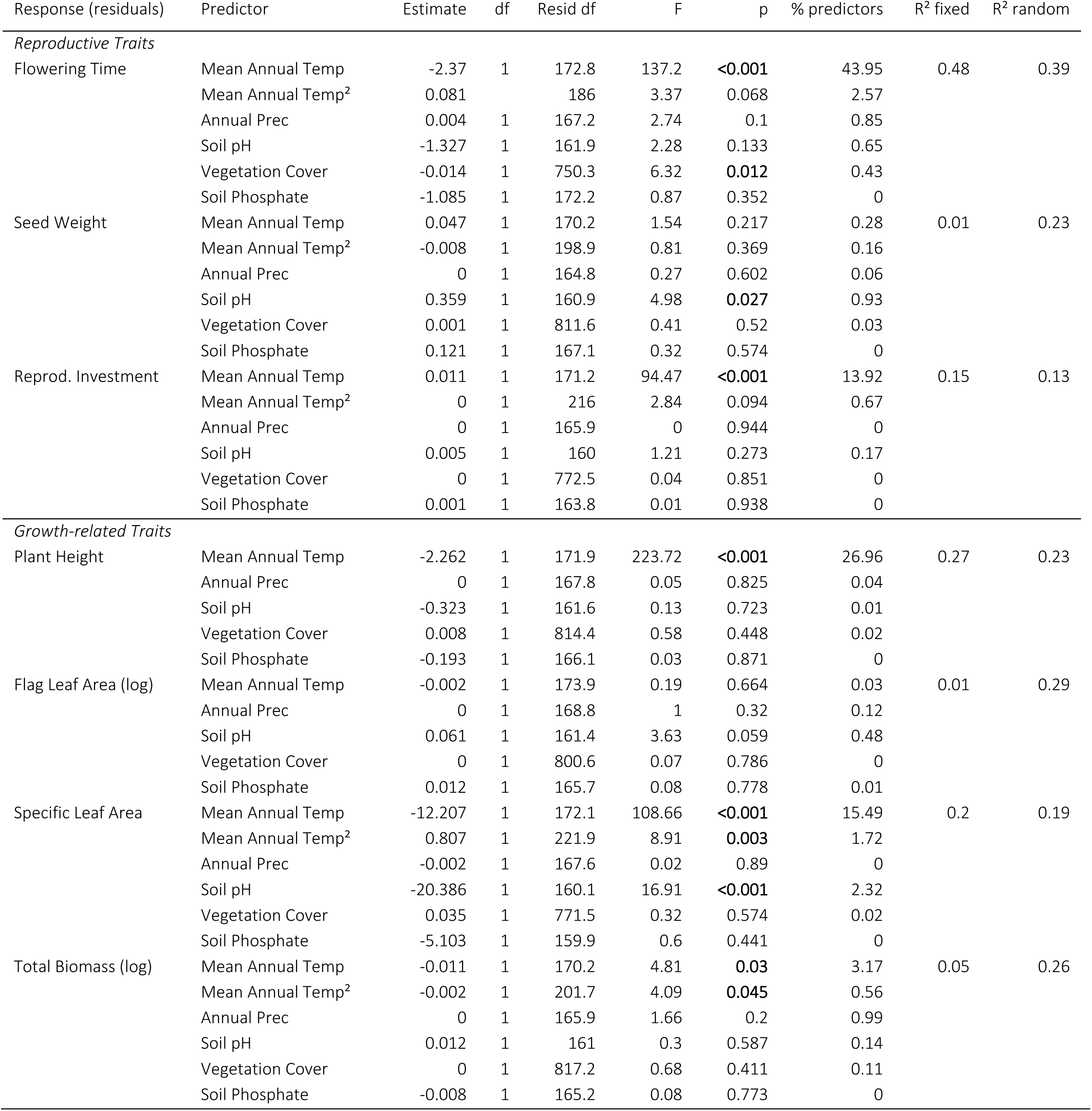
Heritable trait variation in reproductive and growth-related traits testing for the effect of abiotic and biotic factors across large-scale and small-scale gradients. Residuals from a linear model removing fixed treatment effects were modelled against the abiotic and biotic predictors in a linear mixed effect model (LMM), with population and maternal identity as random effects. Mean annual temperature terms, if squared, were centered. Analysis of variance for LMM with error type II. Significant effects (p < 0.05) are highlighted in bold.

**Table S 5.** Estimated fitness optima (peak of the quadratic reaction norm) in relation to mean spring temperature, shown separately for each treatment (Düsseldorf soil, Marburg soil, Marburg sand). Optima were derived from linear mixed effects models fitted with a quadratic temperature term. Confidence intervals were estimated by parametric bootstrapping and 200 simulations (Simulations200x). Historic spring temperature (2000–2020) and 2024 spring temperatures are shown for comparison.

| Treatment | Spring Temperature [°C] |  | Max fitness | CI parametric |  | Simulation200x |  |  |
| --- | --- | --- | --- | --- | --- | --- | --- | --- |
|  | Historic | In 2024 |  | 2.5% | 97.5% | 2.5% | 50% | 97.5% |
| Düsseldorf soil | 12.8 | 13.88 | 14.93 | 14.32 | 15.54 | 14.30 | 14.94 | 15.56 |
| Marburg soil | 10.9 | 12.63 | 13.22 | 8.63 | 14.76 | 7.94 | 13.18 | 14.61 |
| Marburg sand | 10.9 | 12.63 | 15.05 | 14.14 | 16.16 | 14.31 | 15.02 | 16.02 |

